# The Lithuanian reference genome LT1 - a human *de novo* genome assembly with short and long read sequence and Hi-C data

**DOI:** 10.1101/2021.04.05.438426

**Authors:** Hui-Su Kim, Asta Blazyte, Sungwon Jeon, Changhan Yoon, Yeonkyung Kim, Changjae Kim, Dan Bolser, Ji-Hye Ahn, Jeremy S. Edwards, Jong Bhak

## Abstract

We present LT1, the first high-quality human reference genome from the Baltic States. LT1 is a female *de novo* human reference genome assembly constructed using 57× of ultra-long nanopore reads and 47× of short paired-end reads. We also utilized 72 Gb of Hi-C chromosomal mapping data to maximize the assembly’s contiguity and accuracy. LT1’s contig assembly was 2.73 Gbp in length comprising of 4,490 contigs with an N50 value of 13.4 Mbp. After scaffolding with Hi-C data and extensive manual curation, we produced a chromosome-scale assembly with an N50 value of 138 Mbp and 4,699 scaffolds. Our gene prediction quality assessment using BUSCO identify 89.3% of the single-copy orthologous genes included in the benchmarking set. Detailed characterization of LT1 suggested it has 73,744 predicted transcripts, 4.2 million autosomal SNPs, 974,000 short indels, and 12,330 large structural variants. These data are shared as a public resource without any restrictions and can be used as a benchmark for further in-depth genomic analyses of the Baltic populations.

## Background & Summary

The Baltic States consist of three countries, Lithuania, Latvia, and Estonia, located on the eastern coast of the Baltic sea, where Northern, Eastern, and Central European regions converge. The Baltic states share a regional identity^1^ and an endemic LWb blood biomarker is found in high concentrations only in these three populations^2^. It has been established that the Baltic populations were shaped by multiple genetic influxes such as from Anatolia, Mesolithic Western hunter-gatherers, Central Europe^3,4^ as well as a complex history produced by recent wars and annexations. Despite the significance of the Baltic region, the genetic makeup of the Baltic sea region so far has not been studied extensively compared to Central or Southern Europe^3^.

Lithuanians and Latvians have been consistently reported as genetically homogeneous^5–8^ and sharing a very similar genomic structure^8,9^. Until now, genomic research in Lithuanians has mainly utilized single nucleotide polymorphism (SNP) genotyping^5,10–13^ or exome sequencing^14,15^. To expand the scope of analyses and increase the possibility of new findings, whole-genome sequencing (WGS) using long-read technologies is an optimal solution; it enables discovery of novel genomic variations^16^ reveals accurate breakpoints of the structural variations ^4^ and covers some of the complex repeat regions^17–19^. Consequently, resolving haplotypes is also relevant to high quality *de novo* whole-genome assembly and phasing^17^.

Herein, we utilized PromethION, the long-read sequencing platform from Oxford Nanopore Technologies, as a backbone to construct the first Lithuanian reference genome, LT1, using a genome of a healthy female with Lithuanian ancestry. ONT’s PromethION long-read and BGI-500 short-read sequencing technologies were used and merged with Hi-C chromatin conformation capture to complete the genome assembly. Initially two conventional assemblers were used (wtdbg2 ^20^, Shasta ^21^) and better performing one (wtdbg2 ^20^) was selected for the final assembly (polishing and phasing).

The finalized assembly has an N50 value of 138 Mbp and 4,699 scaffolds, and covered 92.75 % of GRCh38, which correspond to a chromosome-scale. Our SV analyses with long-read data identified over twelve thousand consensus SVs, unfortunately, many SV regions could not be annotated indicating that human SVs are an under-investigated area of genomes.

This high-quality assembly is the first step at increasing the availability of human genome assemblies from the Baltic States’ and will serve as a valuable data for further studies in population genomics.

## Methods

### Sample preparation, library construction and sequencing

A Lithuanian female with three generations of ethnic family history was recruited for sequencing. Standard ethical procedures were applied by the Genome Research Foundation with IRB-REC-20101202 – 001. The volunteer signed an informed consent agreement, and a 20ml blood sample was drawn using heparinized needles and collected into anticoagulant containing tubes (K_2_ EDTAA).

DNA was extracted from the donors’ peripheral blood (5 ml) using DNeasy Blood & Tissue Kit from QIAGEN according to the manufacturer’s protocol. Quality and concentration of the extracted DNA were evaluated using NanoDrop^™^ One/OneC UV-Vis Spectrophotometer (Thermo Scientific^™^). Library construction and whole genome sequencing were conducted by Beijing Genomics Institute (BGI) on the BGISEQ-500 platform using the DNBseq^™^ short-read 100bp paired-end sequencing.

Sequencing libraries for long reads were prepared using the 1D ligation sequencing kit (SQK-LSK109) (Oxford Nanopore Technologies, UK) following the manufacturer’s instruction. The products were quantified using the Bioanalyzer 2100 (Agilent, Santa Clara, CA, USA) and the raw signal data were generated on the PromethION R9.4.5 platform (Oxford Nanopore Technologies, UK). Base-calling from the raw signal data was carried out using MinKNOW v19.05.1 with the Flip-Flop hac model (Oxford Nanopore Technologies, UK).

### Hi-C sequencing data generation

Hi-C chromosome conformation capture data were generated using the Arima-HiC kit (A160105 v01, San Diego, CA, USA), and double restriction enzymes were used for the chromatin digestion. To prepare LT1 samples for Hi-C analysis, white blood cells from the donated blood were harvested and cross-linked as instructed by the manufacturer. One million cross-linked cells were used as input in the Hi-C protocol. Briefly, chromatin from cross-linked cells or nuclei was solubilized and then digested using restriction enzymes GATC and GANTC. The digested ends were then labeled using a biotinylated nucleotide, and ends were ligated to create ligation products. Ligation products were purified, fragmented, and selected by size using AMpure XP Beads. Biotinylated fragments were then enriched using Enrichment beads, and Illumina-compatible sequencing libraries were constructed on end repair, dA-tailing, and adaptor ligation using a modified workflow of the Hyper Prep kit (KAPA Biosystems, Inc.). The bead-bound library was then amplified, and amplicons were purified using AMpure XP beads and subjected to deep sequencing. Sequencing of prepared Hi-C libraries was performed using Illumina NovaSeq Platform with read length of 150 bp by Novogene (Beijing, China).

### *De novo* assembly of LT1 genome

For generating the *de novo* assembly of LT1 genome, we prepared a bioinformatic pipeline including: preprocessing step, contig assembly, map assembly, gene prediction, and post analysis. The processes used in the pipeline are summarized in Figure 1.

**Fig. 1.**
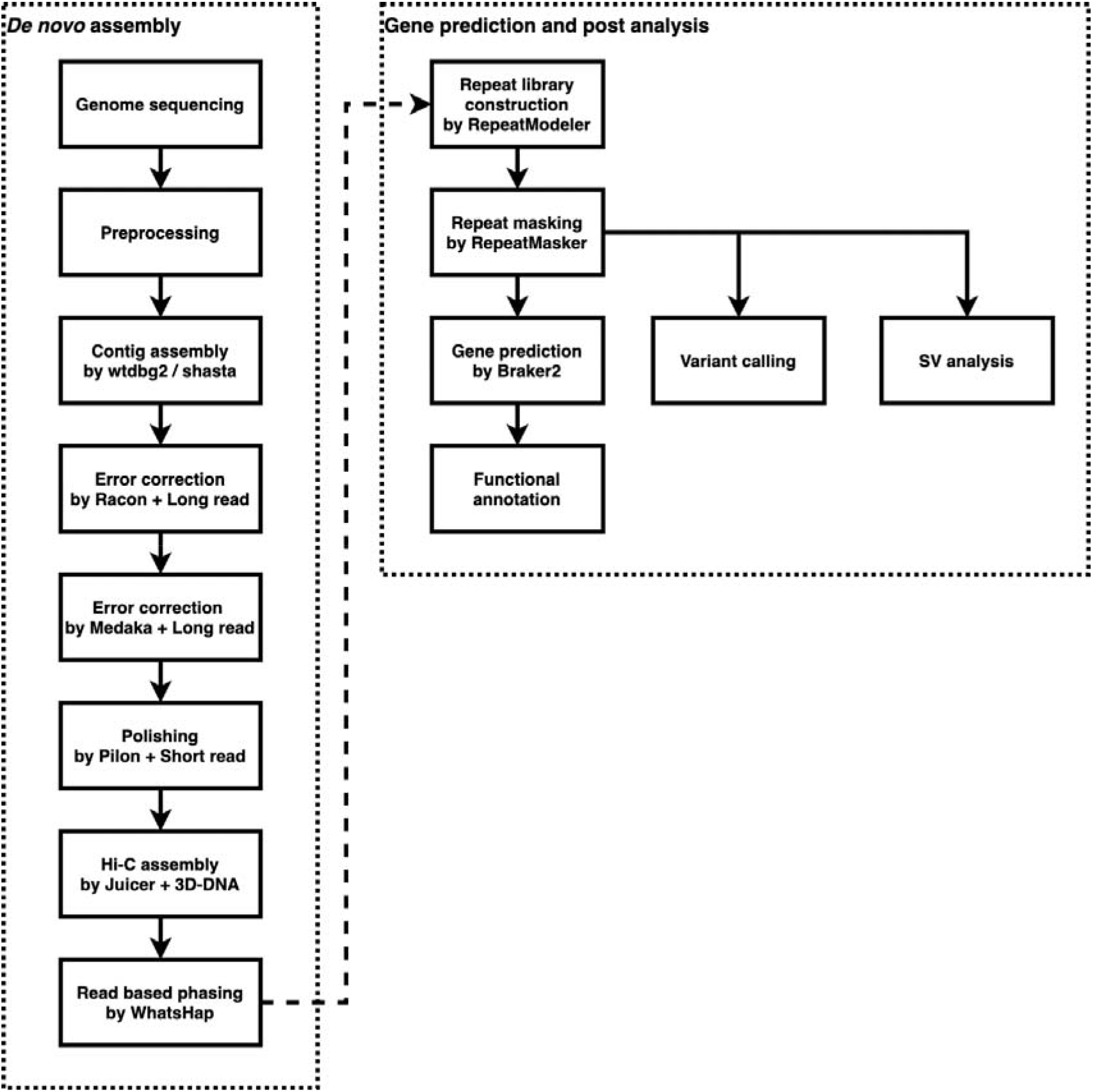
A bioinformatic pipeline for generating LT1 genome assembly.

A total of 142.09 Gbp of short paired-end genomic raw reads were produced by BGISeq 500 sequencer which resulted in a 47×sequencing depth coverage (Table 1). Adapter sequences were trimmed from raw reads using Trimmomatic v0.36^22^ with parameters as ‘ILLUMINACLIP:2:30:10 LEADING:3 TRAILING:3 SLIDINGWINDOW:4:20 HEADCROP: 15 MINLEN:60’, and sequences from vectors and microbial contaminants were removed using BBtools and a customized database from Refseq. For error correction, we used tadpole.sh program of BBtools suite v38.96 (https://sourceforge.net/projects/bbmap). After preprocessing, a total of 106.29 Gbp cleaned reads were obtained.

**Table 1.** Statistics of long and short reads whole genome sequencing for LT1.

| Library type | Sequencing tech. | Library name | Number of reads | Total length of reads (bp) | N50 (bp) |
| --- | --- | --- | --- | --- | --- |
| Long reads | ONT PromethION | LT1_PT | 24,339,507 | 171,726,287,587 | 13283 |
| Short reads | BGISeq-500 | LT1_PE500 | 1,420,906,146 | 142,090,614,600 | 100 |
| Hi-C | Illumina NovaSeq | LT1_HiC | 935,185,202 | 140,277,780,300 | 150 |

A total of 172.22 Gbp raw long reads giving 57× coverage was produced as a result of PromethION sequencing (Table 1). Base-called raw reads with low quality were discarded by the default function of MinKNOW. Adapter sequences were removed using Porechop v.0.2.4 (https://github.com/rrwick/Porechop). After preprocessing, a total of 171.73 Gbp cleaned reads were obtained.

*De novo* assembly was performed using wtdbg2 v2.5^20^ program with cleaned long reads. Parameters for the assembly were set as ‘-x ont -g 3g -L 5000’. For error correction of assembled contigs, we utilized a two-step strategy. First, a consensus generation was carried out using four iterations of Racon v1.4.3 with 4 iterations. The parameters for Racon (https://github.com/isovic/racon) were ‘-m 8 -x -6 -g -8 -w 500’. As the second steps, error correction using Medaka v0.11.5 (https://github.com/nanoporetech/medaka) was performed with a pre-trained model for Flip-flop. To improve the accuracy of the assembly, polishing the consensus sequences with short reads was performed using Pilon v1.23^23^. The polishing step was conducted at two iterations for only SNPs and indels.

Shasta v0.4.0^21^ assembler with the default parameters was used additionally to choose the best assembly from two assemblers. For error correction of assembled contigs, MarginPolish v1.3 (https://github.com/UCSC-nanopore-cgl/MarginPolish) and HELEN v0.0.1 were used with default options^21^.

For generating chromosome-level assembly for the LT1 genome, assembling with 72 Gb of Hi-C reads was performed using Juicer^24^ and 3D-DNA pipeline^25^. Mapping Hi-C DNA reads against assembled contigs were conducted using Juicer. With the mapping information, we proceeded with the 3D-DNA pipeline for LT1 genome. For correcting mis-assemblies of the scaffolds, manual curation was performed using JBAT v1.11.08 (https://github.com/aidenlab/Juicebox/wiki/Juicebox-Assembly-Tools). Assessment for the map assembly was performed using Nucmer v4.00beta2^26^ and Dot (https://github.com/marianattestad/dot) against the human reference genome, GRCh38.

Due to the absence of the LT1’s parental genome data, a read-based phasing of the assembly was performed using Medaka and WhatsHap v1.0^27^ and shared in the LT1 web page. Since the variant calling module of Medaka includes variant calling and phasing steps with sequenced reads from ONT using WhatsHap^27^, cleaned PromethION reads were mapped against the assembled scaffolds and assembled scaffolds were phased using Medaka. As a result of read-based phasing, a total of 2,299,025 variants were phased from whole 3,901,968 variants and number of phased blocks were 8,879 (Table S1). Extracting phased genome sequences was performed using Bcftools v1.9 (http://github.com/samtools/bcftools).

### Construction of repeat library and repeat masking

For repeat masking, we constructed a repeat library for LT1 genome. RepeatModeler v2.0.1 was used with LTRStruct (http://www.repeatmasker.org/RepeatModeler). Repeat masking was conducted using RepeatMasker v4.1.0 (http://www.repeatmasker.org/RepeatMasker).

### Genome annotation

To perform annotation, protein coding genes from GRCh38 were prepared as an evidence gene resource. The gene prediction of LT1 was performed using BRAKER v2.1.4^28^ with GeneMark-ES v4.38^29^ and Augustus v3.3.3^30^. Predicted genes were assessed using BUSCO v4.1.0^31^ with the mammalian orthologous gene set v10. The functional annotation of predicted genes was performed using BLAST+ v2.9.0 against NCBI non-redundant protein ^32^ and Swiss-Prot database^33^.

### Constructing a genome browser and BLAST database

For constructing a genome browser, we compiled all data including predicted gene models and evidence resources. The LT1 genome browser was built using Jbrowse v1.16.9^34^. BLAST database for LT1 gene set v1 was built by SequenceServer v1.0.12^35^.

### Short indel and SNV calling

Variant calling was performed on preprocessed short reads using GATK v4.1.7 HaplotypeCaller^36^ with --output-mode, EMIT_VARIANTS_ONLY and -stand-call-conf 30 settings. Preprocessed short reads were aligned to the GRCh38 reference genome using bwa aligner v0.7.15 (http://bio-bwa.sourceforge.net) and sorting was carried out by samtools v0.1.19^37^. Duplicate marking and quality metric assessment were conducted using picard v1.3.2 (http://broadinstitute.github.io/picard). Base quality scores from the alignment files were recalibrated using BaseRecalibrator and ApplyBQSR tools from GATK^38^. For SNV and indel recalibration dbSNP v146 and Mills_and_1000G_gold_standard.indels sets were used, respectively.

### Structural variation analysis

SVs were identified from pre-processed nanopore reads using sniffles-based meta-pipeline NextSV2^39^ (https://github.com/Nextomics/nextsv) and independent caller SVIM (https://github.com/eldariont/svim). We used a NextSV2^39^ method which employs minimap2, sniffles v1.0.11, and reference GRCh38 with all the default settings for SV calling. For SVIM, default settings were also used. Structural variants with genotypes 0/0 or supported by less than ten reads were filtered out and not presented in the final results. For SV merging and shared variant (as well as union of the variants) estimation, we employed SURVIVOR v1.0.7 (https://github.com/fritzsedlazeck/SURVIVOR) with options 1,000 (bp) for the window size and 30 (bp) for minimum SV length. SURVIVOR merging output was further used for AnnotSV v2.3^40^ multiple database annotation. Copy number variation was estimated using CNVnator v.0.3.3^41^ with default parameters and output filtering settings q0<0.5, where q0 is the fraction of reads mapped with 0 (zero) mapping quality. The complete size distribution of the consensus deletions, insertions, and inversions was plotted in R using packages: data.table (https://cran.r-project.org/web/packages/data.table/index.html), ggplot2^42^, and SiMRiv^43^. Introducing Y axis gaps for deletions plot were created using R package gg.gap (https://rdrr.io/github/ChrisLou-bioinfo/gg.gap/man/gg.gap.html).

## Technical validation

### The statistics of LT1 genome assembly

We assembled long DNA reads produced by ONT PromethION using a blood sample from a female donor. Two conventional assemblers were used: wtdbg2^20^ and Shasta^21^. The contig assembly using wtdbg2 resulted in a total length of 2.73 Gbp, which consists of 4,490 contigs with an N50 of 13.4 Mbp. As for the Shasta assembly, 2.8 Gbp were assembled into 11,009 contigs with an N50 of 6.7 Mbp. Both contig assemblies were corrected in a later stage for errors using long-reads and polished with short-reads as described in methods section. The wtdbg2 assembly had higher contiguity and quality, therefore, it was selected as the main assembly for the LT1 genome and subsequent analyses (Table 2).

**Table 2.** Statistics of contig assembly.

|  | wtdbg2<br>assembly | Shasta<br>assembly | wtdbg2 assembly<br>(> 1kb) | Shasta assembly<br>(> 1kb) |
| --- | --- | --- | --- | --- |
| Contigs No. | 4,490 | 11,009 | 3,659 | 3,573 |
| Total length (bp) | 2,730,857,050 | 2,803,432,513 | 2,730,470,761 | 2,801,466,518 |
| N50 (bp) | 13,411,207 | 6,731,385 | 13,411,207 | 6,731,385 |
| Max contig<br>length (bp) | 65,254,771 | 43,332,451 | 65,254,771 | 43,332,451 |
| Gap | 0.00% | 0.00% | 0.00% | 0.00% |
| GC contents | 40.82% | 40.88% | 40.82% | 40.89% |

The Hi-C data were used for scaffolding the 4,490 contigs. After scaffolding, we acquired 4,699 scaffolds with a total length of 2.73 Gbp and the N50 value of 138 Mbp (Table 3). The number of scaffolds is higher than the original 4,490 contigs because we had to manually split some misassemblies found when we applied the Hi-C data. The longest scaffold was mapped to chromosome 2 and spanned 218 Mbp, which covers 92.6% of chromosome 2. Additionally, using Hi-C long-range mapping information, we were able to scaffold the 4,490 contigs to all 23 chromosomes (Fig. 2). To estimate the quality of the LT1 assembly, we compared it with GRCh3 8^44^ and ‘CHM13 Chromosome X v0.7’ from T2T (https://github.com/nanopore-wgs-consortium/CHM13) using Dot (https://github.com/marianattestad/dot) and NUCmer v3.1^26^. There were no significant misassemblies identified; the LT1 genome covered 92.75 % of GRCh38 (excluding alternative contigs and chromosome Y in the GRCh38), as shown in Fig. 3. Notably, a comparison between LT1 and CHM13 chromosome X displayed a higher concordance (94.50%, Fig. S1), supporting a relatively high-quality assembly of LT1.

**Table 3.** Statistics of final LT1 assembly compared to GRCh38.

|  | LT1 genome v1 | GRCh38 p13 |
| --- | --- | --- |
| Scaffolds No. | 4,699 | 639 |
| Total length (bp) | 2,731,537,123 | 3,272,089,205 |
| N50 (bp) | 138,058,852 | 145,138,636 |
| Max scaffolds length (bp) | 218,797,018 | 248,956,422 |
| Gap | 0.03% | 4.93% |
| GC contents | 40.84% | 41.04% |

**Fig. 2.**
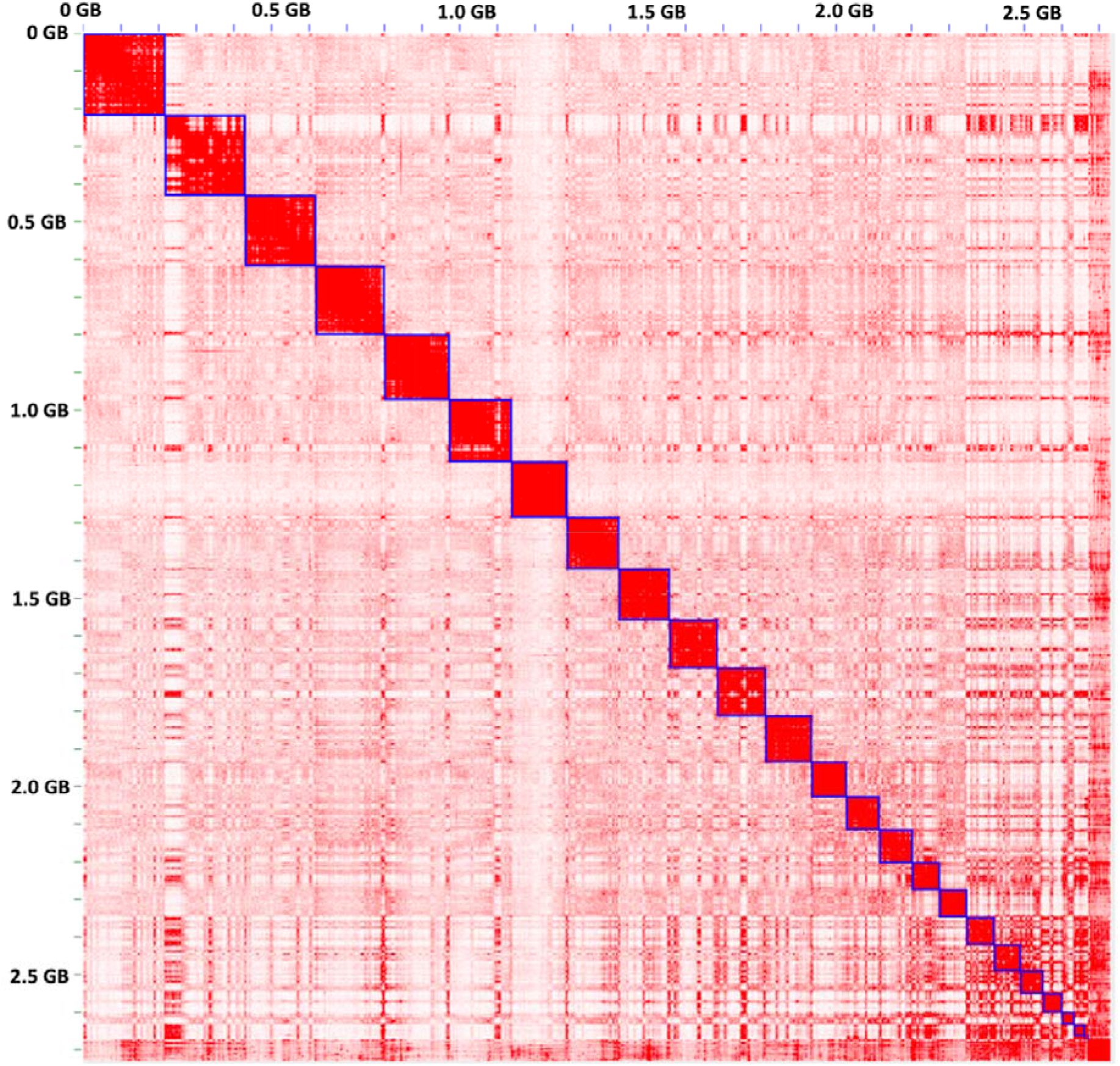
The contact map of Hi-C reads mapped against LT1 assembly.

**Fig. 3.**
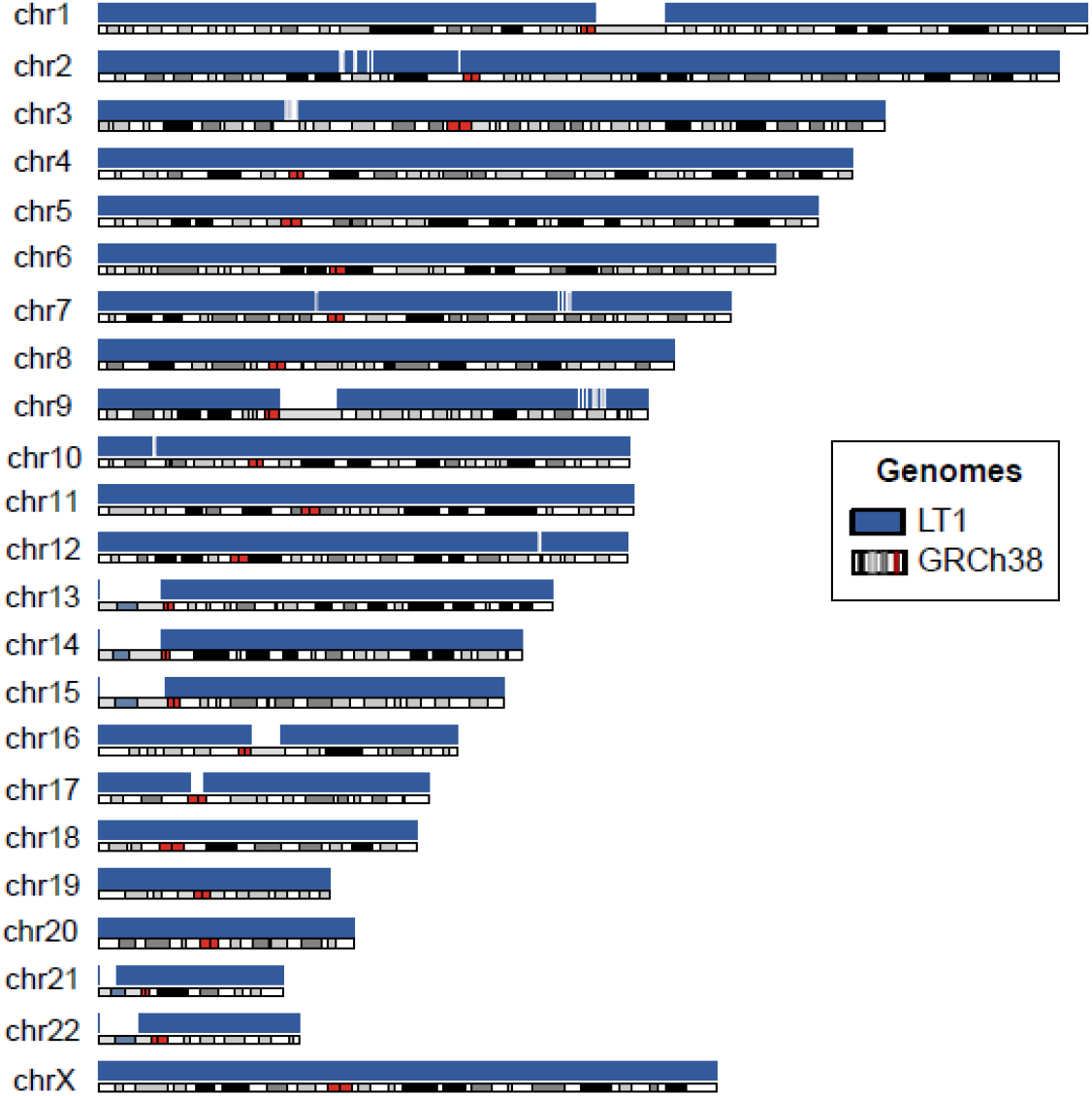
Alignment of LT1 (blue) to the human reference genome GRCh38. The cytobands for the GRCh38 were pre-downloaded from UCSC by the karyoploteR package in R.

The protein-coding gene set in LT1 consisted of 65,614 genes and 73,744 transcripts when we performed gene prediction using BRAKER2^28^. The total length of genes was 86.9 Mbp with an N50 of 1,887 bp. Their GC content was 54.76% and the size of the longest gene, Titin, was 109,026 bp. To assess the gene prediction and assembly quality, we performed a BUSCO^31^ analysis with the mammalian orthologous gene database v10. The ratio of complete single-copy orthologous genes was 89.3% (Table 4). This ratio is slightly below the recommended value, which is likely due to the relatively short assembled genome length (2.73 Gbp) and lack of transcriptome data. The addition of more RNA sequencing data can resolve this incompleteness in the future. The functional annotation using BLAST against the non-redundant protein^32^ and Swiss-Prot database^33^ of the LT1 genome annotated 47,015 (Table S2) and 42,33441,601 transcripts (Table S3), respectively. The result from BLAST analysis is available in http://lithuaniangenome.com.

**Table 4.**
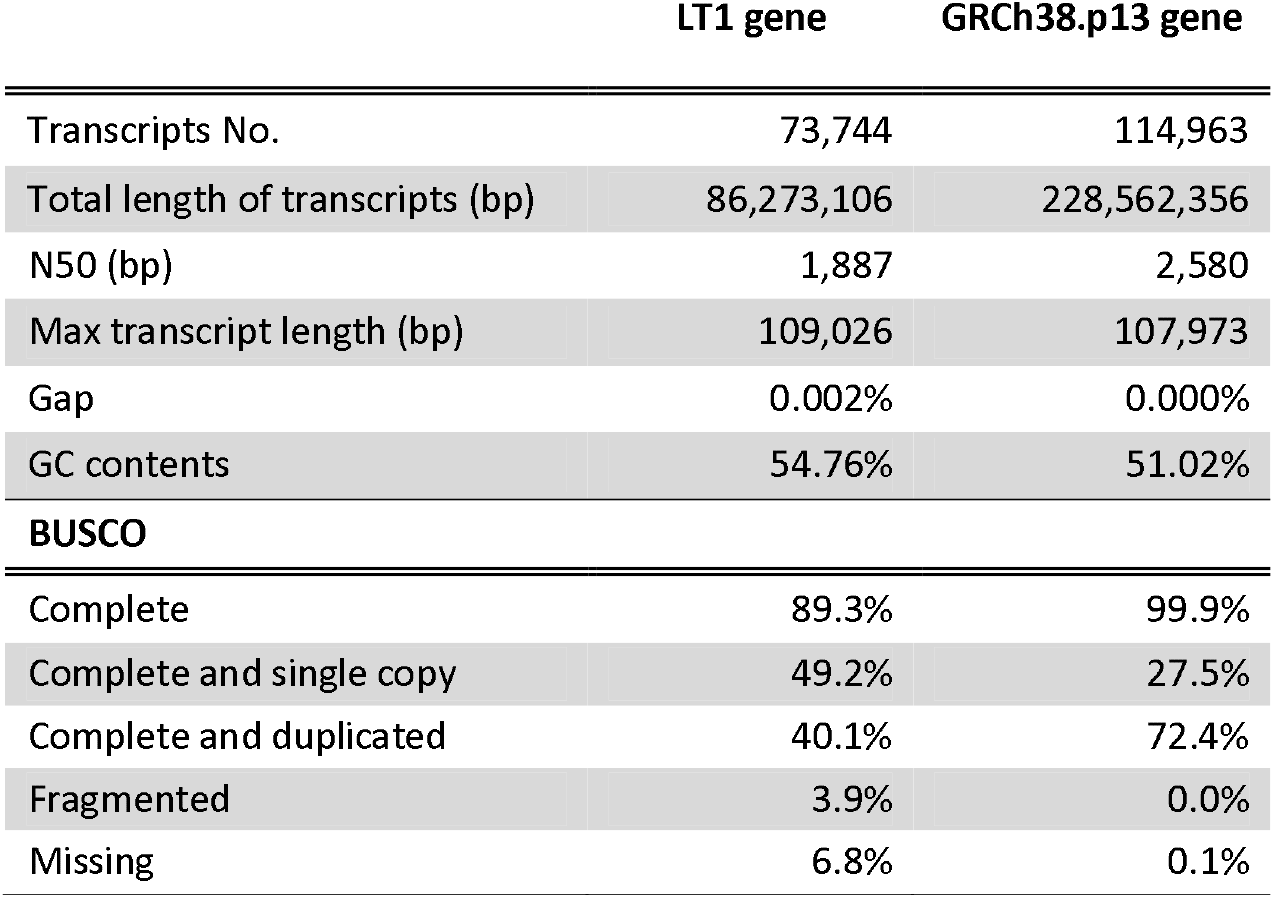
LT1 and GRCh38 genome annotation.

|  | LT1 gene | GRCh38.p13 gene |
| --- | --- | --- |
| Transcripts No. | 73,744 | 114,963 |
| Total length of transcripts (bp) | 86,273,106 | 228,562,356 |
| N50 (bp) | 1,887 | 2,580 |
| Max transcript length (bp) | 109,026 | 107,973 |
| Gap | 0.002% | 0.000% |
| GC contents | 54.76% | 51.02% |
| **BUSCO** |  |  |
| Complete | 89.3% | 99.9% |
| Complete and single copy | 49.2% | 27.5% |
| Complete and duplicated | 40.1% | 72.4% |
| Fragmented | 3.9% | 0.0% |
| Missing | 6.8% | 0.1% |

## Variant identification

LT1 is predicted to have 4,236,954 SNPs and 974,616 indels relative to GRCh38 based on the mapping the short reads against GRCh38. Private variants, variants unreported in dbSNPv.146, are a significant portion of the identified variants (17.81%) (Table S4). Unsurprisingly, heterozygous deletions (30.72%) were the most underreported variant type.

Next, we identified a union of 31,167 SVs, of which, over twelve thousands were consensus insertions, deletions and inversions (Fig. 4, Table S5), using two SV callers, SVIM (https://github.com/eldariont/svim) and NextSV2^39^, and an SV analysis tool SURVIVOR (https://github.com/fritzsedlazeck/SURVIVOR),

**Fig. 4.**
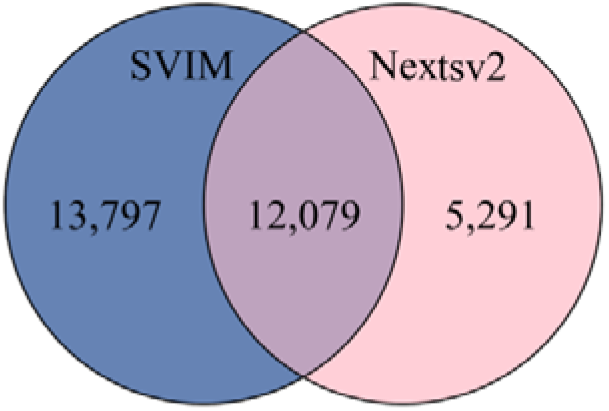
Consensus structural variations found in LT1.

The total number of deletions called by each tool after QC filtering differed minimally, however, the number of insertions identified by SVIM was two times higher than by NextSV2 (Table S5). This is probably because the insertions, in general, are more difficult to call than deletions^45^. CNVnator detected 95 duplications, which was comparable to the 97 by Nextsv2, when minimum read support was 10. Only a small fraction of the SVs could be attributed to a phenotype (10.19%) or found in the major databases such as gnomAD^46^ (37.73%), despite the conservative ten reads threshold to call an SV and confirmation by two different tools. Majority of SVs with phenotypes had a DGV^47^ loss or gain of function annotation (58.21%), (Table S5, Table S6).

Surprisingly, less than half of the consensus SVs could be assigned to a genomic region; almost 97% of the assigned SVs were located in the introns (Table S5). Regardless of the SV type, the small size SVs (30-200 bp length) constituted a significant fraction (64.77%), with another peak spiking around 300 bp (Fig. S2). Among insertions, this region (250-350 bp length) consisted of 81.92% ALU sequences. The longest insertion among the consensus SVs was detected on chromosome 7 in the LOC101928283 gene, spanning 1,003 bases. The largest deletion was in chromosome 1 (45,516 bases) and has already been registered in DGV^47^ and gnomAD^46^ databases despite lacking a precise annotation for location or a clinical phenotype and is predicted to be benign (Table S7).

## Usage notes

We present the first Lithuanian reference genome, LT1. ONT’s PromethION long read and BGI-500 short-read sequencing technologies were used and merged with Hi-C chromatin conformation capture to complete the genome assembly. It was built with enough sequencing data to cover the genome and a high-quality assembly was constructed as the first reference genome from the Baltic States. From the assessment using BUSCO, LT1 gene prediction had more fragmented and missing genes against GRCh38.

Our SV analyses with long-read data showed many regions could not be annotated indicating that SV are an under-investigated area of genomes. Even though the long DNA reads generally provide an advantage of more accurate SV calling, only a small fraction of such variants could be annotated using currently available public databases. More ethnic references and variomes with phenotype association studies are needed to patch these remaining gaps to completely map and understand the biological features of the human genome structure.

## Data Records

The whole genome sequence analyzed in this study has been deposited in the NCBI with a BioProject ID PRJNA635750 and NCBI BioSample database under accession No. SAMN15052346 and in the NCBI SRA database under accession No. PRJNA635750. The LT1 assembly, genome browser, BLAST database and variant calling data can be accessed via http://lithuaniangenome.com.

## Code availability

All the bioinformatic tools used in this project, as well as versions, settings and parameters, have been described in the Methods section. Default parameters were applied if no parameters were specified.

## Acknowledgements

This work was supported by the U-K BRAND Research Fund (1.200108.01) of Ulsan National Institute of Science & Technology (UNIST). This work was also supported by the Research Project Funded by Ulsan City Research Fund (1.200047.01) of Ulsan National Institute of Science & Technology (UNIST). J.B. and Changjae Kim were partially supported by Clinomics Inc. and Genome Research Foundation. We thank GenomeLab, Personal Genomics Institute of Genome Research Foundation, and KOGIC members for providing technical assistance and discussions. We also thank the Korea Institute of Science and Technology Information (KISTI) that provided us with the Korea Research Environment Open NETwork (KREONET).

## Author Contributions

J.B. and A.B. initiated the project as an openfree genome project. H.K., A.B., S.J., Y.K., C.K., C. Y, and J.A. were in charge of methodology, formal analysis and visualization. A.B. and H.K. wrote the manuscript under supervision of J.B. J.B. was in charge of supervision, funding acquisition, and resources. J.B., D.B., A.B., H.K., S.J., C.Y., and J.E. contributed to the manuscript editing process and critical revisions. All authors read and approved the finalized manuscript.

## Ethics declarations

### Approval and participation consent

This study was a part of Korean Personal Genome Project (KPGP also known as PGP-Korea) and was approved by the Institutional Review Board at Genome Research Foundation with IRB-REC-20101202 – 001. The anonymous sample donor has signed a written informed consent to participate in the whole genome sequencing and following analysis in compliance with the Declaration of Helsinki. The (KPGP) informed consent included a section about data publication, which was consented to.

