## Supplementary figures for "The Lithuanian reference genome LT1 - a human *de novo* genome assembly with short and long read sequence and Hi-C data"

**The Lithuanian reference genome LT1 - a human *de novo* genome assembly from the Baltic States**

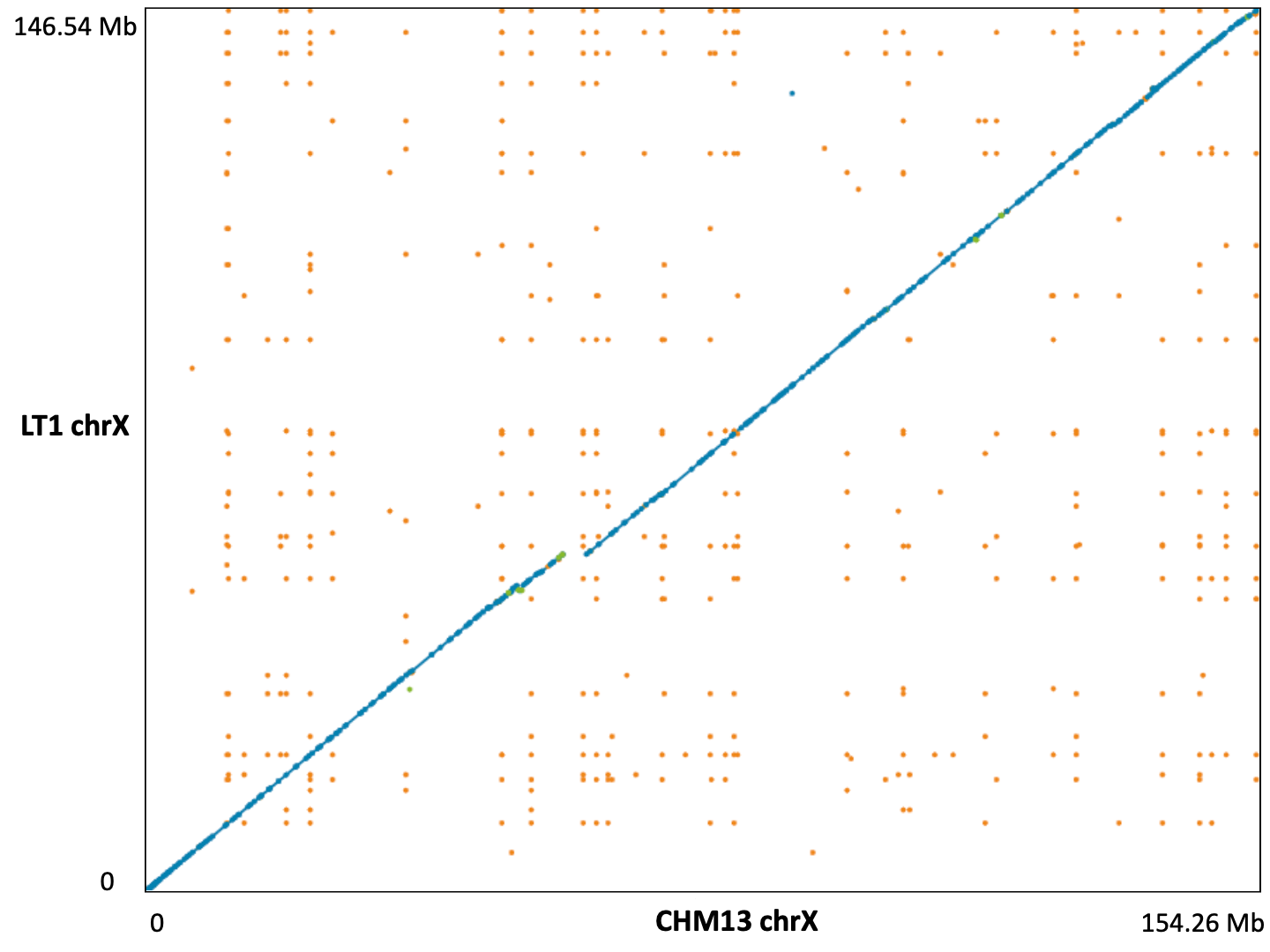

Fig. S1. Comparison of chromosome X between LT1 and CHM13. Blue dots represent unique forward alignments and green dots unique reverse alignments, respectively. Orange dots denote repetitive alignments.

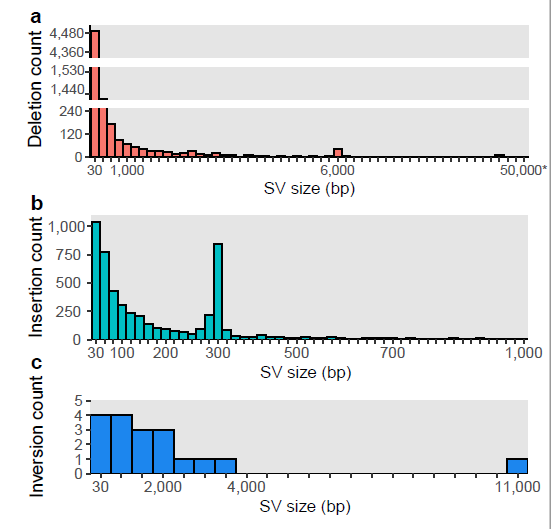

Fig. S2. Complete size distribution of the consensus a) deletions, b) insertions, and c) inversions found in LT1. Number of bins: a) 54, b) 50, c) 21, respectively. Gaps on deletions Y axis are from 250 to 1,198 and from 1,550 to 4,000, aiming to reduce the overrepresentation of the first 2 bins.
